# Evaluation of the role of *whiB6* and *kdpDE* in the dominant multidrug resistant clone *Mycobacterium tuberculosis* B0/W148

**DOI:** 10.1101/2025.01.26.634749

**Authors:** Isabelle Bonnet, Mickael Orgeur, Florence Brossier, Fadel Sayes, Wafa Frigui, Jan Madacki, Hugo Varet, Aurélie Chauffour, Alexandra Aubry, Nicolas Veziris, Wladimir Sougakoff, Roland Brosch, Régis Tournebize

## Abstract

Multidrug resistant (MDR) strains of *Mycobacterium tuberculosis* represent an obstacle to eradicating tuberculosis (TB) due to the low treatment success rate of MDR TB. Among them, the MDR B0/W148 clone has recently evolved from the *M. tuberculosis* Beijing lineage 2 and widely disseminated in Russia and Europe. To get more insights into the genetic factors underlying the evolutionary success of the MDR *M. tuberculosis* B0/W148 clone in addition to environmental and patient-related features, we focused on two mutations specific from this clone that are found in the transcriptional regulators WhiB6 and KdpDE and investigated in a H37Rv strain background the transcriptional profile associated with these mutations and their impact on the *in vitro* and *in vivo* growth characteristics. Through the construction and use of H37RvΔ*whiB6*, H37RvΔ*kdpDE*, and complemented strains, we found that both mutations did not impair the *in vitro* growth of *M. tuberculosis* in standard mycobacterial growth media. The mutation T51P in *whiB6* prevented the upregulation of 9 genes in the *esx-1* core region and 44 genes elsewhere in the genome, while the deletion of two nucleotides in *kdpD* leads to a fusion protein of KdpD with KdpE that inhibits the transcriptional activity of KdpE. Both mutations did not lead to hypervirulence in a mouse infection model. These results point to the role of other MDR B0/W148 specific mutations in the wide geographic diffusion of this clone, and/or put in question a hypothesized hypervirulence as driving factor for this large dissemination.

**Importance:** Human tuberculosis (TB), caused by the bacterium *Mycobacterium tuberculosis,* remains a global public health issue estimated to have been responsible for 1.25 million deaths in 2023. Multidrug resistant (MDR) strains of *M. tuberculosis*, resistant to rifampicin and isoniazid, lead to lower treatment success. Among them, the MDR B0/W148 clone has widely disseminated in Russia and Europe. To get more insights into the genetic factors underlying the evolutionary success of this clone, we investigated two strain-specific mutations found in the transcriptional regulators WhiB6 and KdpDE. By constructing and analysing laboratory *M. tuberculosis* strains carrying these specific mutations, we found numerous changes in their transcriptional profiles, whereas we observed only little impact of these mutations on virulence of *M. tuberculosis* in a mouse infection model. Our study provides new insights into the transcriptional landscape of the selected MDR strains, although no direct connection to virulence could be established.

## Introduction

Human tuberculosis (TB), caused by the etiological agent *Mycobacterium tuberculosis,* remains a global public health issue estimated to have been responsible for 1.25 million deaths in 2023 (1). Multidrug resistance (MDR) of *M. tuberculosis*, *i.e.* resistance to at least isoniazid and rifampicin, represents an obstacle to the WHO objective to reduce TB incidence and deaths as the treatment success rate of MDR TB is 68% compared to 88% when the strain is susceptible (1).

Several reports have described community outbreaks of MDR TB in many areas across the world, with likely contributing factors including variably effective control programs, presence of comorbidities (e.g., HIV co-infection), and numerous patient-related factors, such as societal, immunological and genetic factors (2–4). However, epidemiological studies have highlighted the role of bacterial genetics in the emergence of certain MDR *M. tuberculosis* strains independently of patient and environmental factors (5,6).

Among MDR highly transmissible clones, the B0/W148 MDR Beijing clone, also called Russian clone, has emerged since the early 1960s (5). These strains, belonging to the lineage 2 within the global *M. tuberculosis* phylogeny (7), are the main contributors to the MDR epidemic in Russia and Eastern Europe, and since the USSR’s fall, have also propagated to Western Europe, likely driven by economic or medical migrations of TB patients. According to a study of 720 B0/W148 strains from 23 countries, rifampicin resistance on top of pre-existing isoniazid resistance – and thus MDR – emerged around 1991 and only few strains with a rifampicin susceptibility can be detected nowadays (5,8). In addition to being highly transmissible, the MDR B0/W148 strains seem to be more virulent than non B0/W148 MDR or susceptible Beijing strains and H37Rv, causing a higher bacterial burden *in vitro* in human macrophages and *in vivo* in lungs, spleen and liver of infected mice (9,10). Moreover, a Russian clinical study including 144 cases of tuberculous spondylitis showed a 2-fold increase in the rate of Beijing B0/W148 strains in the spinal TB group versus the pulmonary TB group, also suggesting a greater virulence of this clone (11). However, the underlying mechanisms explaining this B0/W148 evolutionnary success remain unknown.

Here we investigated mutations of representative 100-32 *M. tuberculosis* strains isolated in France, 100-32 being the main 24-locus mycobacterial interspersed repetitive unit-variable number tandem repeat (MIRU-VNTR) code of B0/W148, which might have contributed to the evolutionary success of B0/W148 strains (5). Among the various mutations that were identified as being specific for the MDR B0/W148 clone, we focused on two potentially involved in virulence, found in the transcriptional regulators KdpDE and WhiB6. We characterized the transcriptional profiles associated with these mutations in the H37Rv *M. tuberculosis* model strain background and evaluated their potential impact on the *in vitro* and *in vivo* growth characteristics of *M. tuberculosis* strains.

## Results

### Comparative genomic analysis reveals two specific mutations in *whiB6* and *kdpDE*

To identify single nucleotide polymorphisms (SNPs) or small insertions and deletions (indels) specific to the MDR B0/W148 clone, we probed genome sequences of *M. tuberculosis* strains available at the National Reference Centre of Mycobacteria. Our collection comprised 32 isolates that showed a 24-locus MIRU-VNTR genotype named MLVA Mtbc 15-9 “100-32”, of which 3 were non MDR. Traditionally, the Beijing B0/W148 type was defined using *IS6110*-RFLP typing whereas 100-32 corresponded to the main MIRU-VNTR code of it (12). In order to increase the specificity, we compared the genomes of 29 MDR 100-32 strains with genomes of the 3 non MDR 100-32 isolates. Further comparison with 457 other isolates, mostly MDR, representing *M. tuberculosis* lineages 1, 2, 3 and 4, identified 30 non-synonymous SNPs and small deletions specific to the MDR 100-32 clone (Table 1). Among them, we decided to investigate the impact of the mutations identified and previously reported as specific variants of the MDR B0/W148 clone in *whiB6* (*Rv3862c*) and *kdpD* (*Rv1028c*) (8,13). Indeed, both genes are transcriptional regulators described to be implicated in *M. tuberculosis* virulence (14,15). These genes are located at two different genomic loci in *M. tuberculosis*, with *whiB6* located upstream of the core region of the ESX-1 type VII secretion system, and *kdpD* upstream of the *kdpFABC* operon at genome coordinates 4338 kb and 1151-1149 kb (reverse strand) of strain H37Rv, respectively.

**Table 1.**
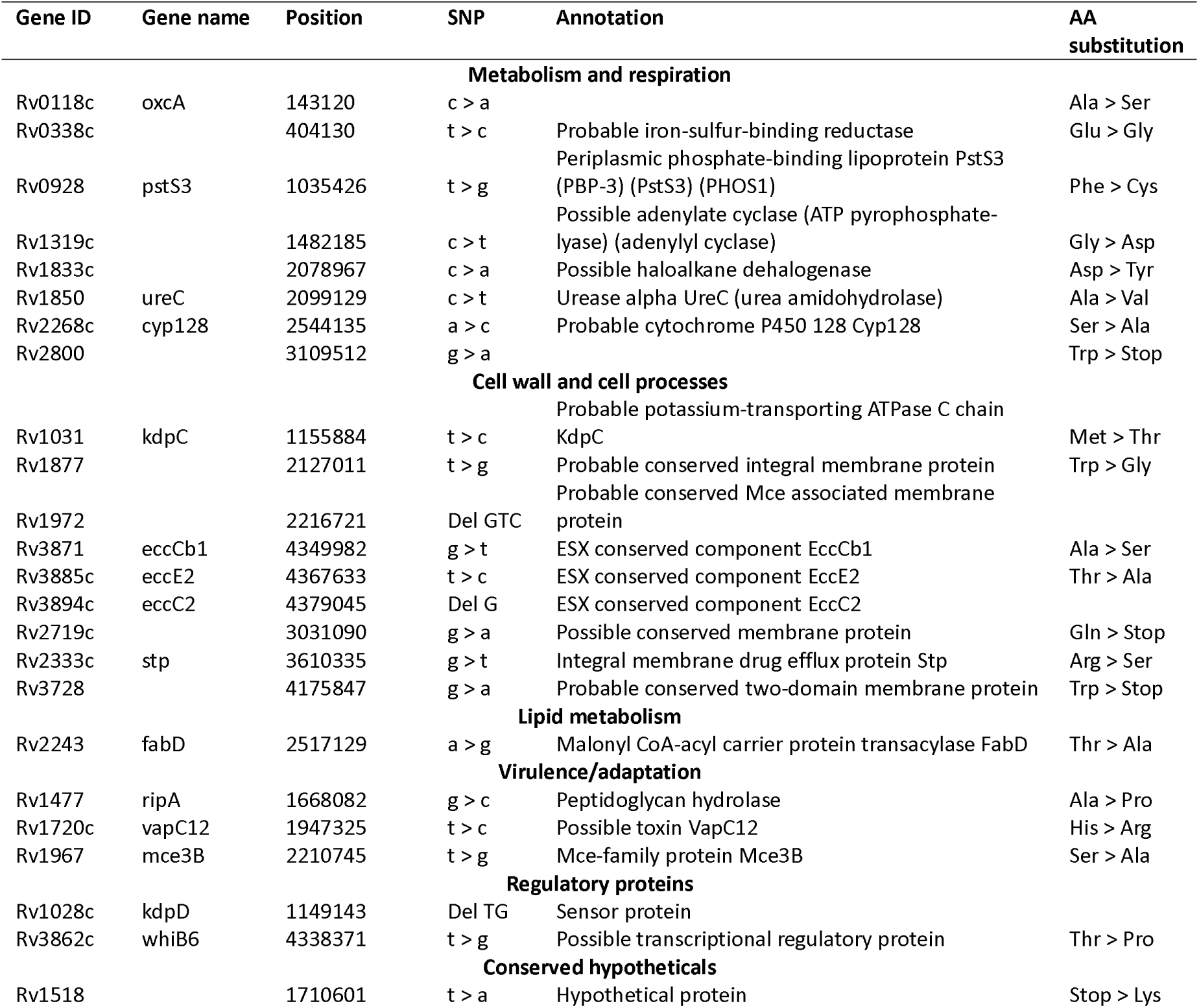
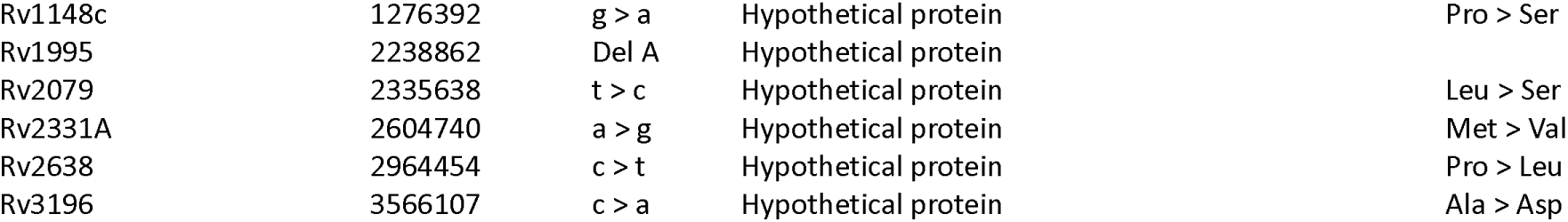
Non-synonymous mutations and small deletions specific to the MDR *M. tuberculosis* 100-32 clone. Positions are given relative to H37Rv reference sequence (NC_000962.3). Small deletions are given in 5’-3’ direction. Abbreviations: SNP, single nucleotide polymorphism; AA, amino acid.

The non-synonymous mutation in *whiB6* replaces threonine with proline at position 51 in WhiB6. Previous work showed that an insertion of a unique single nucleotide G in the *whiB6* promoter region in both H37Rv and H37Ra reference strains relative to clinical strains uncoupled the ESX-1 system essential for *M. tuberculosis* pathogenesis from its regulation via WhiB6 as part of the PhoP regulon (14).

The *kdpD* and *kdpE* genes code for a sensor with histidine kinase activity and a transcription factor, respectively. Both genes are located upstream of the *kdpFABC* operon coding for a potassium transport system (16). A KdpDE deletion mutant strain of *M. tuberculosis* showed higher virulence in a mouse model (15). In MDR B0/W148, the deletion in *kdpD* of two nucleotides (c.2541_2542delCA, hereafter referred as ΔCA) downstream of the region coding for the essential phosphorylation site (His642) leads to a fusion of the sensor KdpD to the regulator KdpE.

In order to investigate the impact of the MDR B0/W148 specific *whiB6* T51P mutation and the *kdpDE* ΔCA deletion, we deleted these genes in the H37Rv reference strain background (H37RvΔ*whiB6* and H37RvΔ*kdpDE*), and complemented them with various WT and mutant constructs. We used the H37Rv reference strain model for easier handling and security reasons, thereby avoiding the manipulation of large culture volumes of MDR strains. The Δ*whiB6* mutant strain was complemented with 3 constructs: *i*) promoter G insertion - *whiB6* WT, corresponding to the organisation in H37Rv and H37Ra; *ii*) promoter WT - *whiB6* WT, corresponding to the organisation in clinical isolates; and *iii*) promoter WT - *whiB6* T51P, corresponding to the organisation in MDR B0/W148 strains (Fig. 1A). The Δ*kdpDE* mutant strain was complemented by either the WT or the MDR B0/W148-specific ΔCA sequences of *kdpDE* (Fig. 1B).

**Fig. 1.**
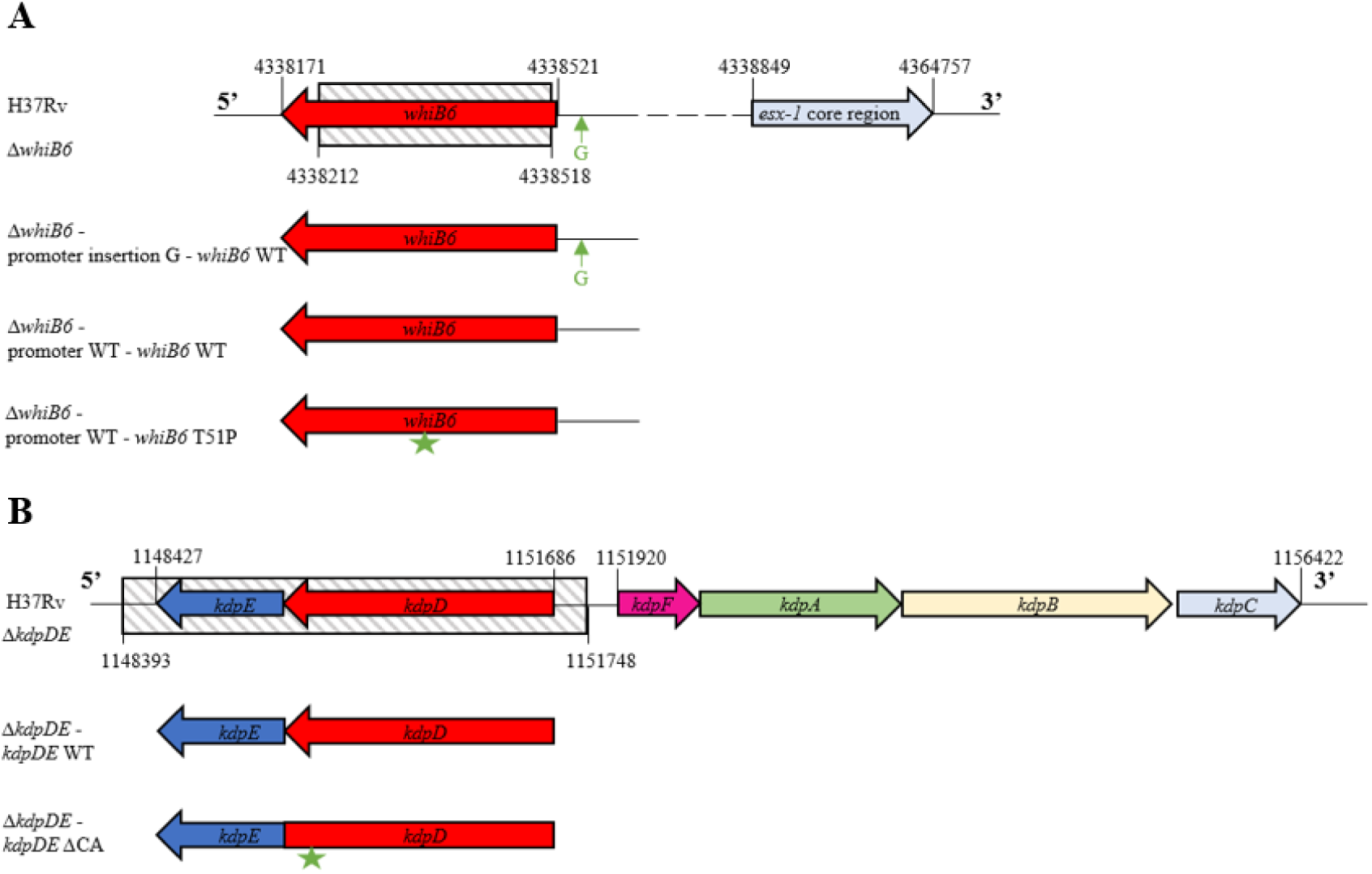
Schematic representation of the genomic organisation of *whiB6* and *kdpDE* in *M. tuberculosis* H37Rv. (A) Δ*whiB6* mutant and complemented strains. (B) Δ*kdpDE* mutant and complemented strains. Hatched area corresponds to the region deleted in the mutant strain.

### The mutation T51P in *whiB6* and the deletion in *kdpDE* do not impact the *in vitro* growth

To check the impact of the mutations on bacteria fitness, we measured the kinetics of *in vitro* growth of the various strains in different media. In 7H9 broth, we did not observe a significant difference between the growth of the H37Rv strain, the Δ*whiB6* mutant strain and any of the mutant strains complemented with either *whiB6* WT or *whiB6* T51P (Fig. 2A), indicating that the *whiB6* T51P mutation does not affect *in vitro* growth. As KdpDE act as sensor and regulator of K^+^ uptake, we measured the growth rate in 7H9 medium containing either 7 mM K^+^ (normal concentration) or without K^+^. H37Rv, Δ*kdpDE*, and WT or ΔCA-complemented strains grew identically in medium with 7 mM K^+^ or without K^+^, although they grew more slowly in medium without K^+^ (Fig. 2B,C). These results show that the MDR B0/W148 specific mutations in *whiB6* and *kdpDE* do not alter the normal growth rate and fitness of the bacteria *in vitro* in standard mycobacterial growth media.

**Fig. 2.**
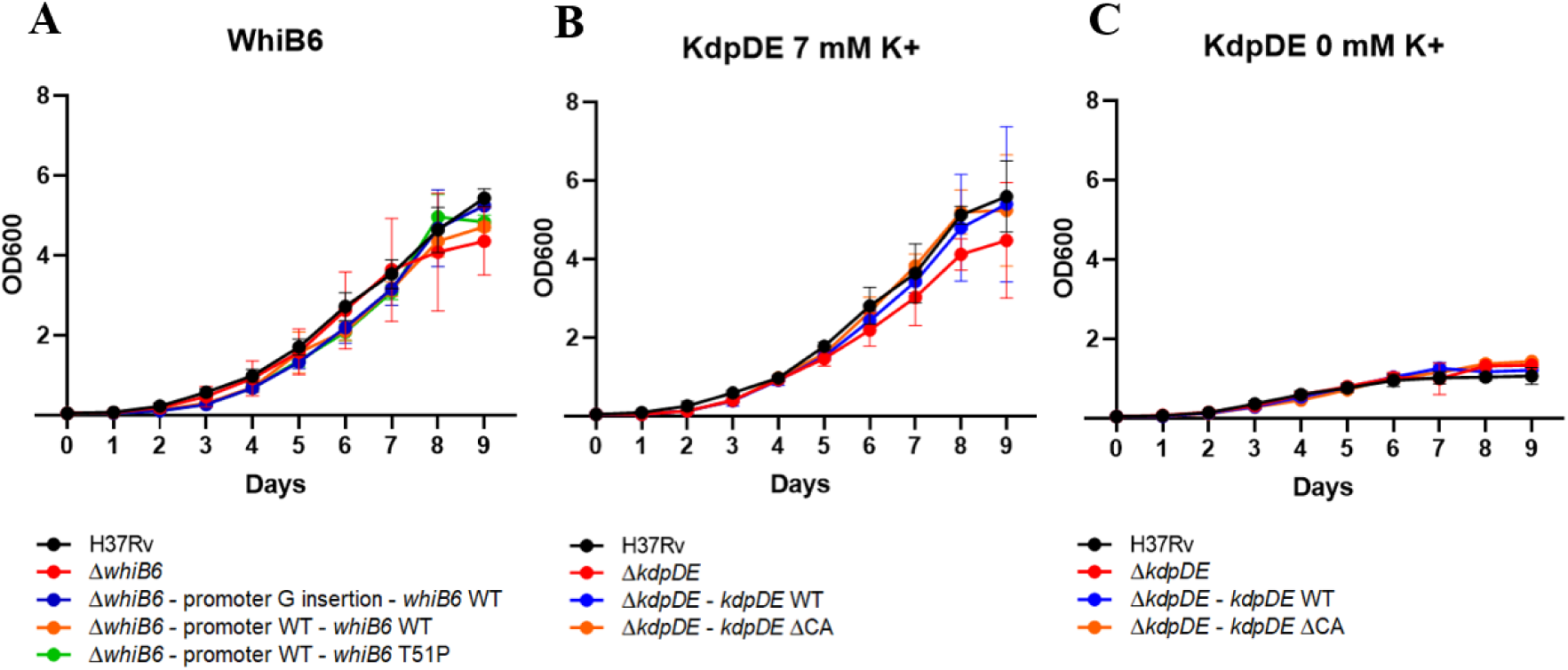
Impact of the MDR B0/W148 specific mutations in *whiB6* and *kdpDE* on the *in vitro* growth rates of H37Rv WT, *whiB6* and *kdpDE* mutant and complemented strains. (A) Growth rate of WT, Δ*whiB6* mutant and complemented strains in standard medium. (B, C) Growth rate of WT, Δ*kdpDE* and complemented strains in normal (B) and potassium-depleted (C) conditions. Results are from three independent experiments and are expressed as the mean optical density (OD) ± SD at 600 nm.

### The mutation T51P in *whiB6* impacts the expression of a large geneset, including genes of the core *esx-1* locus

To get more insights into the effect of the *whiB6* T51P mutation on the transcriptional role of WhiB6, we analyzed and compared the transcriptional landscape of H37Rv, the Δ*whiB6* mutant strain, the promoter WT - *whiB6* WT complemented strain, and the promoter WT - *whiB6* T51P complemented strain. We identified a total of 339 genes that exhibited significant changes in expression compared to H37Rv (Fig. 3A, Table S1, Fig. S1), with 191 (56.3%) of those genes that were specific to the Δ*whiB6* mutant strain or to the promoter WT - *whiB6* WT complemented strain (Fig. 3B, Table S1). Among those genes, 9 genes located in the core region of the ESX-1 type VII secretion system were upregulated significantly in the strain carrying the promoter WT - *whiB6* WT complementation construct relative to the H37Rv strain, while expression of those genes was unchanged in the Δ*whiB6* mutant strain or the promoter WT - *whiB6* T51P complemented strain, except for *Rv3872* (*pe35*), which was downregulated significantly in the Δ*whiB6* mutant strain (Fig. 3A,C). These results are consistent with a previous study reporting that the mutation in the *whiB6* promoter region, present in strains H37Rv and H37Ra, impairs the transcriptional activation of the ESX-1 system via WhiB6 (14). They also indicate that the *whiB6* T51P mutation has a detrimental effect on the functionality of WhiB6 in the H37Rv background, resulting in a lack of transcriptional activation of the *esx-1* locus similar to the Δ*whiB6* mutant strain. To confirm this, we analyzed the expression and secretion of EsxA (ESAT-6) and EsxB (CFP-10) in *M. tuberculosis* strains H37Rv, CDC1551, H37RvΔRD1, H37RvΔ*whiB6*, and complemented variants. The CDC1551 strain is a highly transmissible clinical isolate (17), whereas the H37RvΔRD1 strain is an attenuated H37Rv strain lacking part of the ESX-1 secretion system (18). We observed a strong expression of EsxA and EsxB in the whole cell lysate (WCL) of bacteria, as well as a strong secretion in the supernatant (SN) fraction for the strain CDC1551 (Fig. 3D). By contrast, both EsxA and EsxB expression and secretion were drastically reduced in H37Rv, Δ*whiB6*, the promoter G insertion - *whiB6* WT and the promoter WT - *whiB6* T51P complemented strains. Only the complementation with the WT *whiB6* under the expression of the active promoter (promoter WT - *whiB6* WT) induced a strong expression and secretion of both EsxA and EsxB, similar to what is observed with the CDC1551 strain. These results show that the T51P mutation drastically impairs WhiB6 transcriptional regulation function in the H37Rv background leading to reduced expression of EsxA and EsxB. The transcriptional regulation alteration of WhiB6 T51P was also noticeable at its transcript level since the promoter WT - *whiB6* T51P complemented strain depicted a higher *whiB6* expression abundance as compared to the promoter WT - *whiB6* WT complemented strain (Fig. 3E), thus suggesting an auto-regulation of *whiB6* via a negative feedback loop.

**Fig. 3.**
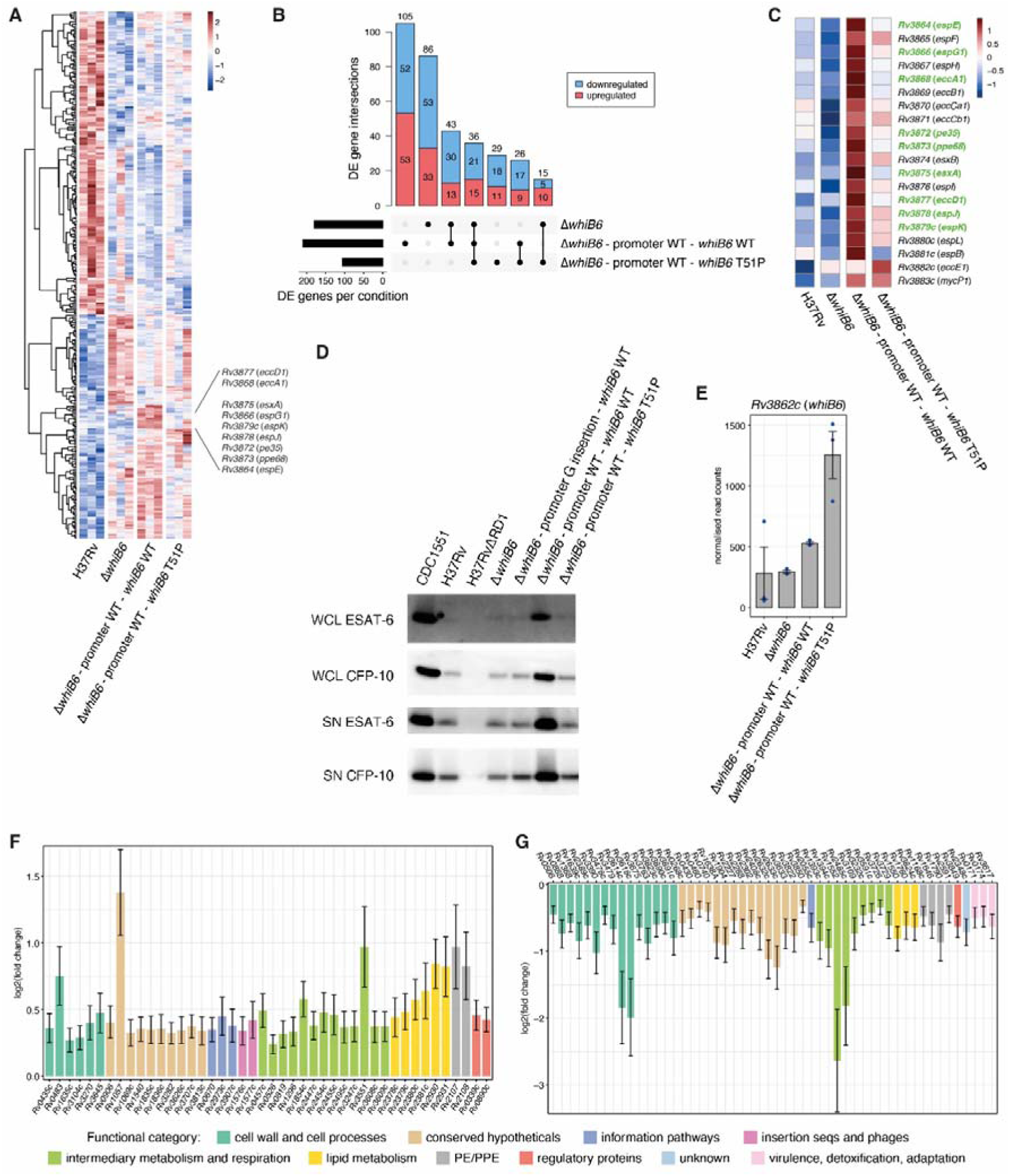
Transcriptional effects of the MDR B0/W148 mutation T51P in *whiB6* on gene expression during exponential growth. (A) Hierarchical clustering of the 339 differentially expressed genes detected across the Δ*whiB6* mutant strain, the promoter WT - *whiB6* WT complemented strain, and the promoter WT - *whiB6* T51P complemented strain as compared to H37Rv. Gene expression levels were scaled by row across all genotypes and replicates (colorbar). (B) Intersection of differentially expressed genes detected in each condition (black bars). Black circles in the bottom matrix layout indicate the genotypes that are part of each intersection. Number of genes within each intersection set is indicated at the top of each bar. Downregulated genes are depicted in blue while upregulated genes are depicted in red. (C) Relative expression levels of the genes belonging to the core *esx-1* locus across all genotypes. Gene expression levels were averaged across all replicates per genotype and scaled by row (colorbar). Genes that were significantly detected as differentially expressed in the promoter WT - *whiB6* WT complemented strain are highlighted in green. (D) EsxA (ESAT-6) and EsxB (CFP-10) production (WCL, Whole Cell Lysate) and secretion (SN, supernatant) in CDC1551, H37Rv, H37RvΔRD1, *whiB6* mutant or complemented strains. This experiment was done in triplicate per strain. (E) Relative transcript abundances of *whiB6* gene in each genotype. Blue circles represent the normalized read counts determined in each replicate. Bars and error bars represent the mean ± s.e.m. (F,G) Expression changes of genes detected as significantly upregulated (F) and downregulated (G) specifically in the promoter WT - *whiB6* WT complemented strain relative to H37Rv. Color of the bar indicates the functional category the genes belong to.

To further investigate the transcriptional changes induced in the promoter WT - *whiB6* WT complemented strain, we also focused on the 96 genes located outside of the core *esx-1* locus that were detected as differentially expressed specifically in this genotype. Given that their expression was unchanged in the Δ*whiB6* mutant strain or the promoter WT - *whiB6* T51P complemented strain, these genes are potentially activated (for those upregulated) or repressed (for those downregulated) by functional WhiB6 but not by T51P WhiB6. Regarding the upregulated genes, 44 belonged, for the most represented categories, to the categories “intermediary metabolism and respiration” (13 genes), “conserved hypotheticals” (10 genes), “lipid metabolism” (6 genes), and “cell wall and cell processes” (6 genes) (Fig. 3F, Table S1). In regard to lipid metabolism, 2 of the 6 genes, named *fadD26* (*Rv2930*) and *ppsA* (*Rv2931*), encode enzymes that are involved in the synthesis of phthiocerol dimycocerosate (PDIM), a well-established virulence lipid of *M. tuberculosis* (19–22). By contrast, 52 genes were downregulated specifically in the strain carrying the promoter WT - *whiB6* WT complementation construct. These genes were assigned, for the most represented categories, to the categories “cell wall and cell processes” (15 genes), “conserved hypotheticals” (15 genes), and “intermediary metabolism and respiration” (9 genes) (Fig. 3G, Table S1). Among genes annotated as being involved in cell wall and cell processes, *espA* (*Rv3616c*) and *espD* (*Rv3614c*) from the *espACD* operon and *espR* (*Rv3849*, annotated as ESX-1-related regulatory protein) were downregulated unlike the genes of the ESX-1 core region (Fig. 3C,F, Table S1), potentially leading to a balanced regulation of the ESX-1 system. Indeed, EspR activates the expression of the *espACD* operon (23,24), whose expression promotes EspA and EspC co-dependent secretion of EsxB and EsxA (25,26). Taken together, our results show that the direct and/or indirect regulation of WhiB6 on gene expression in *M. tuberculosis* is much wider than the previously reported effects on genes of the ESX-1 core region and thereby open new insights into the regulatory network of this important member of the WhiB regulatory family.

### The ΔCA deletion in *kdpD* inhibits the activation of the *kdpFABC* locus upon potassium depletion

We next investigated the effect of the MDR B0/W148-specific *kdpDE* ΔCA deletion and determined the transcriptome of H37Rv, the Δ*kdpDE* mutant strain and the *kdpDE* ΔCA complemented strain grown in 7H9 culture medium with or without potassium. Given that the strongest effect on transcript abundances was associated with the presence or absence of potassium, differential expression analysis was performed by comparing the Δ*kdpDE* mutant strain and the *kdpDE* ΔCA complemented strain against their respective H37Rv control in each growth condition separately. In normal growth with potassium, 300 genes exhibited significant changes in expression in the Δ*kdpDE* mutant strain or the *kdpDE* ΔCA complemented strain as compared to H37Rv (Fig. 4A,B, Table S2, Fig. S2), whereas upon potassium depletion, 679 genes were differentially regulated (Fig. 4C,D, Table S3, Fig. S3). In contrast to H37Rv, for which *kdpD* and *kdpE* genes were upregulated upon potassium depletion, the expression of both genes was already activated in the *kdpDE* ΔCA complemented strain in the presence of potassium (Fig. 4A,C,E). As a consequence, the *kdpFABC* locus was upregulated during normal growth condition in the *kdpDE* ΔCA complemented strain at an expression level that was similar to the one in the Δ*kdpDE* mutant strain (Fig. 4E, Table S2), and that was mostly unchanged upon potassium depletion contrary to H37Rv (Fig. 4E, Table S3). These results thus suggest that the MDR B0/W148-specific ΔCA mutation-mediated fusion of proteins KdpD and KdpE either impairs the sensor activity of KdpD to properly activate KdpE, and/or prevents the transcriptional activity of KdpE to upregulate the *kdpFABC* operon in response to the absence of potassium.

**Fig. 4.**
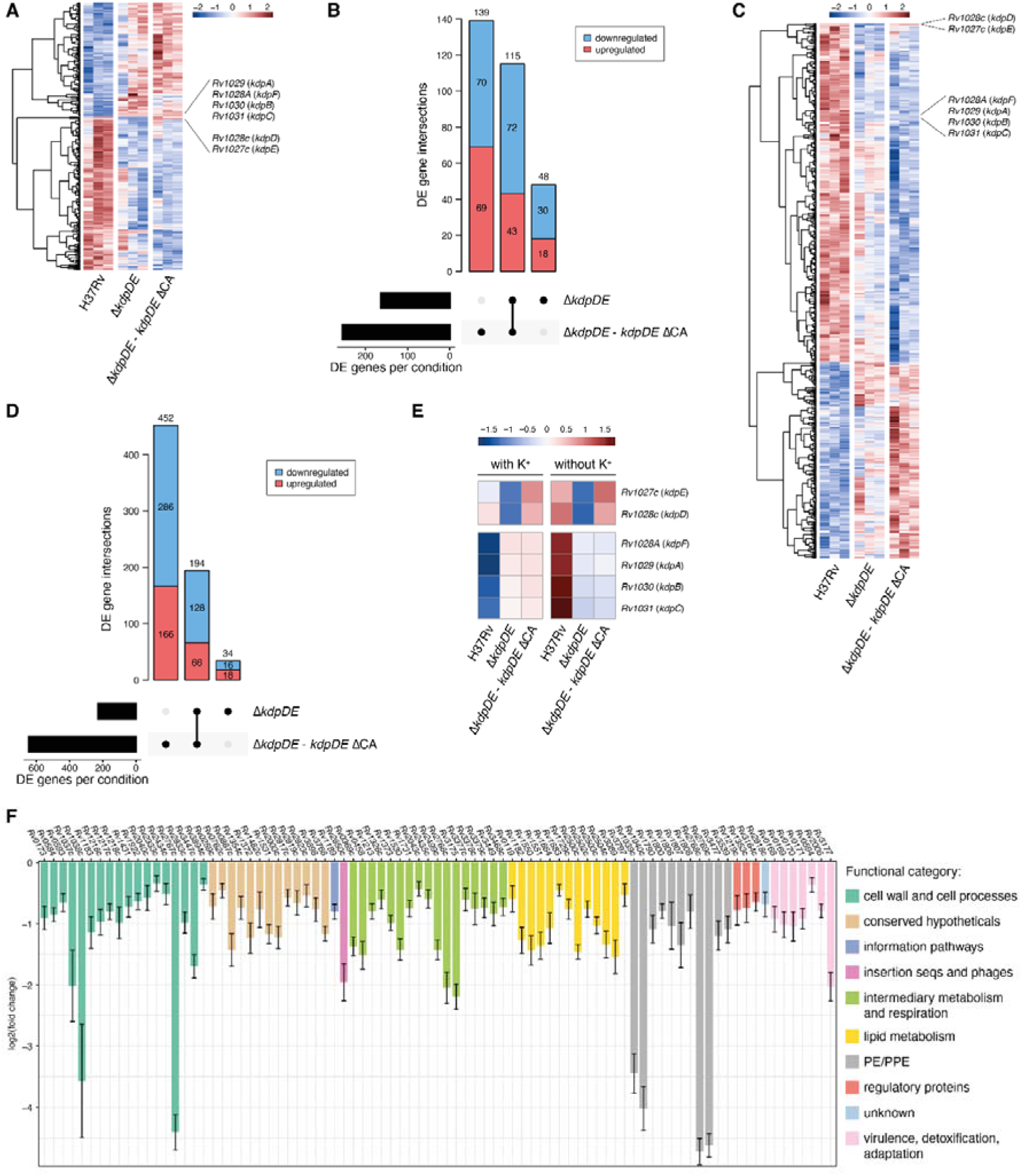
Transcriptional effects of the MDR B0/W148. ΔCA deletion in *kdpD* on gene expression during exponential growth with and without potassium. (A) Hierarchical clustering of the 300 differentially expressed genes detected in the Δ*kdpDE* mutant strain and the *kdpDE* ΔCA complemented strain as compared to H37Rv during growth with potassium. Gene expression levels were scaled by row across all genotypes and replicates (colorbar). (B) Intersection of differentially expressed genes detected in each genotype with the presence of potassium (black bars). Black circles in the bottom matrix layout indicate the genotypes that are part of each intersection. Number of genes within each intersection set is indicated at the top of each bar. Downregulated genes are depicted in blue while upregulated genes are depicted in red. (C) Hierarchical clustering of the 679 differentially expressed genes detected in the Δ*kdpDE* mutant strain and the *kdpDE* ΔCA complemented strain as compared to H37Rv during growth without potassium. Gene expression levels were scaled by row across all genotypes and replicates (colorbar). (D) Intersection of differentially expressed genes detected in each genotype with the absence of potassium (black bars). Black circles in the bottom matrix layout indicate the genotypes that are part of each intersection. Number of genes within each intersection set is indicated at the top of each bar. Downregulated genes are depicted in blue while upregulated genes are depicted in red. (E) Relative transcript abundances at the *kdpDE* and *kdpFABC* loci between genotypes and growth conditions. Gene expression levels were averaged across all replicates per condition and scaled by row (colorbar). (F) Expression changes of downregulated genes relative to H37Rv in the *kdpDE* ΔCA complemented strain that were shared with the Δ*kdpDE* mutant strain and specifically detected in the growth condition without potassium. Color of the bar indicates the functional category the genes belong to.

As the *kdpDE* operon is known to be activated in low potassium concentrations, we further investigated the genes that were differentially expressed only in this particular growth condition. Upon potassium depletion, 128 genes were significantly downregulated in both Δ*kdpDE* mutant strain and *kdpDE* ΔCA complemented strain as compared to H37Rv (Fig. 4D, Table S3). Among these genes, 85 were specifically downregulated only in the absence of potassium, and thus correspond to genes normally activated by KdpE in response to potassium shortage. The most represented functional categories among these 85 genes were “cell wall and cell processes” (18 genes), “intermediary metabolism and respiration” (17 genes), and “lipid metabolism” (13 genes) (Fig. 4F).

Among the 17 genes involved in intermediary metabolism and respiration, 7 encode enzymes that belong to the class of oxidoreductases, possibly involved against the oxidative stress encountered during infection. Apart from oxidative stress, GabD2 (*Rv1731*) is one of these oxidoreductases, which forms an alternative pathway from alpha-ketoglutarate to succinate, along with GabD1 and Kgd, in the tricarboxylic acid cycle (27). In addition, 3 genes encode enzymes involved in glucose metabolism.

Concerning lipid metabolism, 3 downregulated genes code for FadD enzymes: FadD7 (*Rv0119*), FadD11 (*Rv1550*), and FadD13 (*Rv3089*). In *M. tuberculosis*, there are 34 fatty acid adenylating enzymes (FadD) that can be grouped into two classes: fatty acyl-CoA ligases (FACLs) involved in lipid and cholesterol catabolism; and long chain fatty acyl-AMP ligases (FAALs) involved in the biosynthesis of numerous essential and virulence-conferring lipids found in *M. tuberculosis*. FadD7, FadD11, and FadD13 are part of FACLs, which convert free fatty acids into acyl-coenzyme A thioesters, the first step in fatty acid degradation (28). Additionally, two downregulated genes encode the FadE enzymes FadE19 (*Rv2500c*) and FadE35 (*Rv3797*). These enzymes are acyl-CoA dehydrogenases, which introduce unsaturation into fatty acids (29). Most of the FadE enzymes characterized to date in *M. tuberculosis* function in cholesterol catabolism and play roles in the dehydrogenation of cholesterol substrates through β-oxidation.

### *whiB6* T51P and *kdpDE* ΔCA do not increase virulence in a mouse infection model

Next we assessed the role of *whiB6* T51P and *kdpDE* ΔCA mutation in *M. tuberculosis* ability to grow and persist *in vivo* 30 and 60 days after BALB/c mice infection with H37Rv, the Δ*whiB6* and Δ*kdpDE* mutant strains, and the various complemented strains. As shown in Fig. 5A, the log10 fold changes in the bacterial load at day 30 or 60 relative to the initial inoculum measured at day 1 (Fig. S4) for the Δ*whiB6* mutant strain, the promoter G insertion - *whiB6* WT, the promoter WT - *whiB6* WT, and the promoter WT - *whiB6* T51P complemented strains, show that the bacteria were able to survive and grow in the lungs. At day 30 post-infection, the growth of the promoter G insertion - *whiB6* WT complemented strain was statistically lower than that of the promoter WT - *whiB6* WT complemented strain. At day 60, the growth of the promoter G insertion - *whiB6* WT complemented strain reached a level similar to the other strains. The bacterial load of the promoter WT – *whiB6* T51P complemented strain was significantly lower than that of H37Rv or the Δ*whiB6* mutant strain. Taken together, the virulence evaluation of the Δ*whiB6* mutant and complemented strains revealed overall quite similar bacterial growth for all tested strains. These findings suggest that WhiB6 is not a key player for modulating mycobacterial virulence in mice, and that in a similar fashion the WhiB6 T51P mutation in the MDR B0/W148 strains might only have very limited impact, if any, on the virulence level of this strain.

**Fig. 5.**
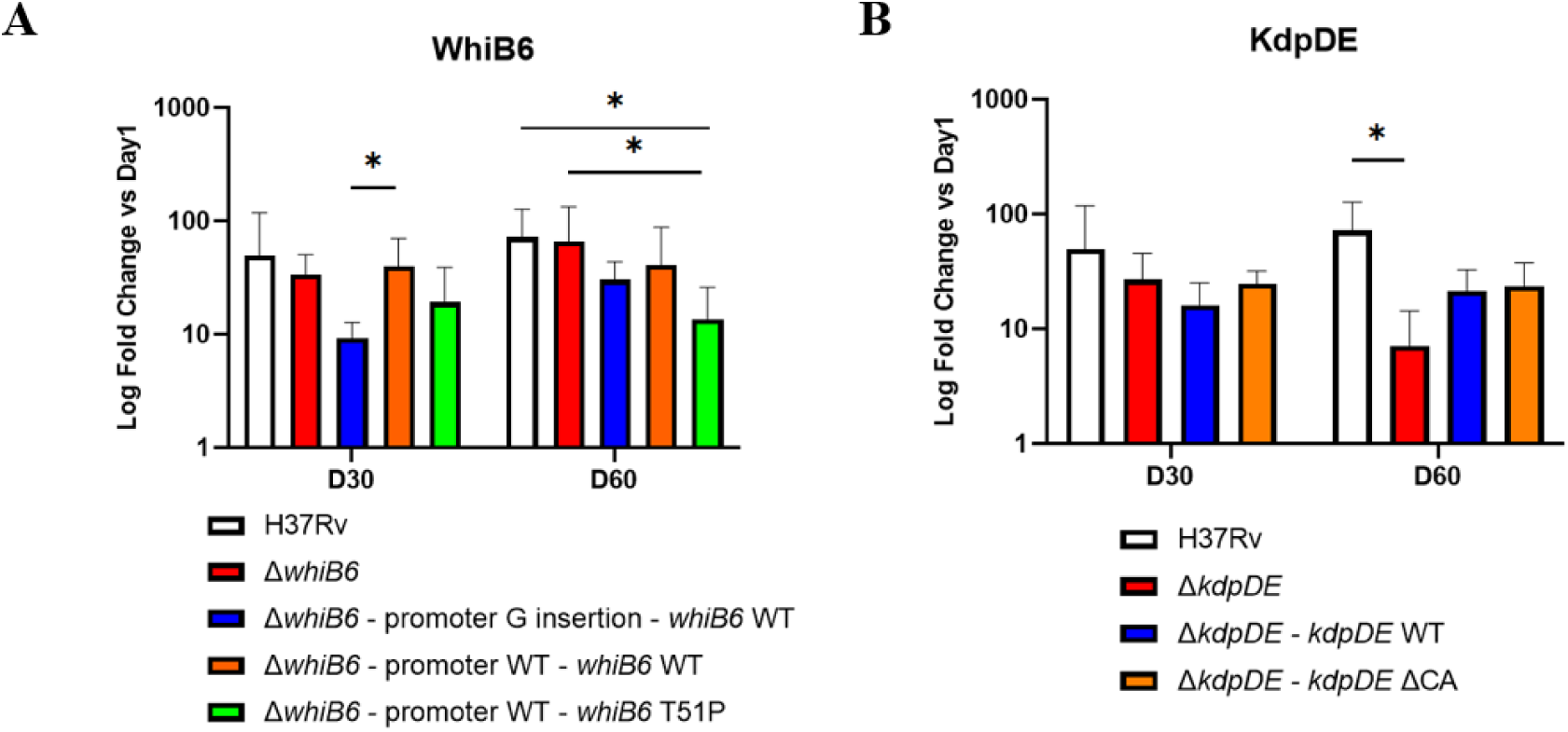
Virulence evaluation of *whiB6* T51P and *kdpDE* ΔCA mutations in BALB/c mice. (A) Time course after mouse infection with H37Rv, the Δ*whiB6* mutant strain, and the promoter G insertion - *whiB6* WT, the promoter WT - *whiB6* WT, and the promoter WT - *whiB6* T51P complemented strains. (B) Time course after mouse infection with H37Rv, the Δ*kdpDE* mutant strain, and both *kdpDE* WT and ΔCA complemented strains. Values represent fold change in the lung bacterial load at day 30 or day 60 relative to day 1. Means are for 5 to 10 mice per condition.

The growths of the Δ*kdpDE* mutant strain, and both *kdpDE* WT and ΔCA complemented strains were found equivalent at day 30 post-infection (Fig. 5B). In contrast, at day 60, the number of CFU recovered from lungs of mice infected with the Δ*kdpDE* mutant strain was lower than that recovered from lungs of mice infected with H37Rv but did not significantly differ from the *kdpDE* WT or ΔCA complemented strains. The log10 fold changes in mice infected with the *kdpDE* WT complemented strain or with the *kdpDE* ΔCA complemented strain were identical at day 60. We conclude from these results that the Δ*kdpDE* mutant strain had an *in vivo* growth disadvantage at day 60 relative to the H37Rv WT strain that was only partially complemented by the *kdpDE* or *kdpDE* ΔCA constructs. While we do not know the reasons for this only partial complementation, especially for the *kdpDE* WT version, our results from the 60 days timepoint suggest that KdpDE plays some role in virulence regulation in *M. tuberculosis*.

## Discussion

MDR TB outbreaks represent a brake on WHO’s End TB Strategy. In this context, the much wider diffusion of some MDR clones is of strong concern. Our goal was to characterize genetic modifications of strains belonging to the MDR B0/W148 clone that might explain its great diffusion, in addition to environmental or patient-related factors. We included 32 sequenced 100-32 clinical isolates of our strains collection, 100-32 being the main MIRU-VNTR code of B0/W148 strains (12) : 29 were MDR and 3 non MDR. When compared with the 3 non MDR strains but also with 457 other, mostly MDR, isolates, representing *M. tuberculosis* lineages 1, 2, 3 and 4, we identified in the 29 MDR strains 30 non-synonymous mutations and small deletions specific to the MDR 100-32 clone and sought to investigate in the H37Rv genetic background the role of two mutations found in the *whiB6* and *kdpDE* regulatory genes, which were previously reported as specific variants of the MDR B0/W148 clone in other studies (8,13).

We were particularly interested in KdpD as it is part of the two-component system KdpDE, which controls, in response to environmental stimuli, the expression of the inducible, high-affinity K^+^ transporter KdpFABC (16,30,31). In the MDR B0/W148 clone, the deletion of two nucleotides (ΔCA) at the end of *kdpD* leads to a fusion protein KdpDE that could impact the regulatory function of KdpE by restricting its localisation to the membrane. In our study, no significant growth difference was noted in medium with (7 mM) or without (0 mM) potassium between H37Rv, the Δ*kdpDE* mutant strain, and its *kdpDE* WT and ΔCA complemented strains, suggesting that the activity of these genes are not essential for *in vitro* growth. For *kdpE*, this observation differs from an initial high density transposon mutation analysis in which *kdpE* was found to to be required for optimal *in vitro* growth (32), whereas in a later study, a few *kdpE* transposon insertion mutant strains could be observed (33). Additionally, we observed a lower growth of H37Rv, the Δ*kdpDE* mutant strain, and the complemented strains in medium without K^+^, as described in a model of dormant state (34). We here describe that *kdpDE* under the conditions we used were not essential for *in vitro* growth of *M. tuberculosis*. This observation opens new perspectives on the essentiality information availability for selected genes in the *M. tuberculosis* genome, which apparently may vary depending on the *in vitro* growth conditions and/or the type of genetic modification. Our transcriptomic analysis of H37Rv, the Δ*kdpDE* mutant strain, and the *kdpDE* ΔCA complemented strain during exponential *in vitro* growth revealed that the absence of *kdpDE* or the MDR B0/W148 version of ΔCA in *kdpDE*, in the presence of potassium, resulted in a constitutive expression of *kdpFABC* with higher expression of these genes compared to H37Rv, suggesting that KdpE normally represses the expression of *kdpFABC* when potassium is not lacking. By contrast, reduced expression of *kdpFABC* was seen in Δ*kdpDE* and *kdpDE* ΔCA complemented strains upon depletion of potassium relative to H37Rv as *kdpFABC* was more highly upregulated by phospho-KdpE in H37Rv than in both mutant and complemented strains, which remained at its basal expression level between the presence and absence of potassium. These results indicate that the MDR B0/W148 ΔCA mutation in *kdpDE* impairs the transcriptional activity of KdpE at two levels in the H37Rv background. It not only prevents the repression of *kdpFABC* when potassium is available but also affects the upregulation of *kdpFABC* when potassium is limited.

Although the deletion of *kdpDE* or the ΔCA mutation in *kdpDE* neither affected the *in vitro* growth nor induced major changes on the global transcriptional level, we observed a reduced replication of the Δ*kdpDE* mutant strain at a late time point in BALB/c mice compared to the H37Rv WT strain (Fig. 5B), while complementation with both *kdpDE* WT and *kdpDE* ΔCA revealed only partial complementation of the attenuation of Δ*kdpDE* mutant strain. However, our results are in stark contrast to the results published for the 1691-bp *kdpDE* deletion mutant strain that showed enhanced virulence in SCID mice in previous work (15). Such discordant observations could be explained by the differences in the mouse model, such as the use of immune-deficient versus immune-competent mice, and the fact that only survival rates were measured in the previous study, which may show different outcomes compared to results based on CFU. In any case, even it was previously suggested that inactivation of *kdpDE* leads to hypervirulence of *M. tuberculosis* in SCID mice (15), which was one of the arguments why we had chosen to investigate the *kpdDE* ΔCA mutation of MDR B0/W148, the combined results of our study suggest that the ΔCA-mediated fusion of *kpdD* and *kdpE,* corresponding to a loss of function mutation, does not cause hypervirulence of *M. tuberculosis*, at least in H37Rv, as could have been hypothesized based on this previous report (15).

Our results also show that the transcriptional function of WhiB6 is impaired by the T51P mutation in a H37Rv background. This protein encoded at the 5’ end of the *esx-1* core region belongs to the WhiB family of proteins, found exclusively in Actinobacteria (35). In the MDR B0/W148 clone, a non-synonymous mutation in *whiB6* replaces threonine with proline at position 51. It was therefore of interest to evaluate if this mutation possibly enhanced or reduced the regulation activity of WhiB6 in this emerging strain family. We did not observe an impact of this mutation or the gene deletion on *in vitro* growth, suggesting that it did not impair the fitness. This is consistent with previous work indicating that *whiB6* is not an essential gene for H37Rv *in vitro* growth (32). Growth was also not affected by the sequence of *whiB6* promoter (with or without a G at the −74 position relative to the *whiB6* start codon). Our transcriptomic analysis of *M. tuberculosis* H37Rv, the Δ*whiB6* mutant strain and the complemented strains revealed 339 differentially regulated genes, including 9 from the *esx-1* core region. While the transcription level of these genes in the promoter WT - *whiB6* WT was upregulated, the transcriptional level in the promoter WT - *whiB6* T51P complemented strain was much lower and similar to the one of H37Rv, indicating that the T51P mutation impairs the function of WhiB6. Interestingly, many other genes outside the *esx-1* core region were also differentially regulated and thus potentially subject to WhiB6-mediated regulation. A striking example was the downregulation of the EspR regulator and the *espACD* cluster in the promoter WT - *whiB6* WT complemented strain, suggesting that WhiB6 may also fulfill repressor functions, contributing to a fine-tuned secretion of ESX-1 substrates. When considering the regulation of the 9 genes from the *esx-1* core locus, the results are further substantiated by the lower *in vitro* production and secretion of the EsxA and EsxB proteins in the promoter WT - *whiB6* T51P complemented strain compared to the clinical strain CDC1551, which has been involved in a TB outbreak (17). This low level of EsxA and EsxB similar to the one observed in H37Rv, which is known to express *esxA* and *esxB* at basal levels, did not impact the ability of *M. tuberculosis* to replicate in BALB/c mice, a finding that is in agreement with the well-established virulence phenotype of strain H37Rv in mouse infection models.

Our present work revealed many new details on *M. tuberculosis whiB6* and *kpdDE* function and their regulatory networks, and in some part confirms previous work, but also generates new questions on the biological impact of these two important regulatory components of *M. tuberculosis*. However, as these two mutations, which are preserved in all the MDR B0/W148 isolates, did not show a substantial impact on the bacterial fitness under *in vitro* or *in vivo* conditions as conducted in a H37Rv-based *M. tuberculosis* model, it is also likely that some of the other 28 MDR B0/W148 non-synonymous mutations and small deletions, alone or in combination, might be more impactful. We originally hypothesized that some of the mutations might create hypervirulence, but found that this was not the case in our model system for the *whiB6* and *kdpDE* mutations. Our observations may also indicate that hypervirulence might not be the explanation of the wide diffusion of the MDR B0/W148 clone. Indeed, some mutations in clinical strains, such as in *kdpD* (36,37), have been well associated with clustering. For instance, a different host immune response, rather than hypervirulence, could result in a higher transmissibility of the clone (38). By characterizing high and low transmission strains of *M. tuberculosis* in mice, it has been shown that high transmission *M. tuberculosis* strain induces granulomas with the potential to develop into cavitary lesions that aids bacterial escape into the airways and promotes transmission. It remains for the moment unknown what genetic changes might be behind the emergence and wide distribution of the MDR B0/W148 clone, but it is likely that the question is more complex than previously thought, likely involving the interplay of several different non-synonymous mutations or indels in addition to environmental traits, thereby opening new and interesting perspectives to better understand the emergence of specific *M. tuberculosis* strain families and lineages.

## Materials and methods

### Bacterial strains and culture conditions

*E. coli* DH10B and DH5a strains, used for cloning procedures, were grown on LB agar medium and/or LB broth. *M. tuberculosis* strains were obtained from stock held at the French National Reference Centre for Mycobacteria or Institut Pasteur (for CDC1551 and H37RvΔRD1 strains). Mycobacterial strains were cultured in Middlebrook 7H9 broth supplemented with 10% OADC (oleic acid, albumin, dextrose, and catalase, BD) and 0.05% Tween 80 or on Middlebrook 7H11 medium supplemented with 10% OADC. When required, antibiotics were included for selection purposes at the following concentrations: ampicillin (100 µg/mL), hygromycin (150 µg/mL), kanamycin (20 µg/mL), and zeocin (25 µg/mL) for *E. coli*; hygromycin (50 µg/mL), kanamycin (20 µg/mL), and zeocin (25 µg/mL) for mycobacteria.

For potassium limitation studies, *M. tuberculosis* strains were cultured in Middlebrook 7H9 broth supplemented with 10% OADC and 0.05% Tween 80 containing either 7 mM or 0 mM K^+^ in which case KH_2_PO_4_ was replaced by NaH_2_PO_4_.

### Mutant strain generation and complementation

The *M. tuberculosis* H37Rv mutant strains were constructed by allelic replacement using the recombineering method (39). The allelic exchange substrate *whiB6*::zeo was obtained by a three step PCR approach (40). Briefly, two 500-bp fragments corresponding to the *whiB6* upstream and downstream regions were amplified by PCR from the *M. tuberculosis* H37Rv genomic DNA and linked to a third PCR fragment encoding the zeomycin resistance cassette, to generate the 1.6 kb-fragment *whiB6*::zeo. The NEBuilder HiFi DNA Assembly Cloning Kit (Biolabs) was used to obtain the allelic exchange substrate *kdpDE*::zeo. The *whiB6*::zeo and *kdpDE*::zeo fragments were thus used to transform a *M. tuberculosis* H37Rv recombinant strain containing the pJV53 vector. The pJV53 plasmid encodes the recombination proteins gp60 and gp61 (41), whose expression is induced at mid-logarithmic phase by incubation with 0.2% acetamide for 24h (39). The putative mutant strains were selected on solid medium for resistance to kanamycin and zeomycin. Deletion of *whiB6* and *kdpDE* was confirmed by whole genome sequencing. Spontaneous loss of the pJV53 plasmid was obtained by serial rounds of culture without kanamycin.

Five different complementation integrative pYUB412-based plasmids harbouring the *whiB6* and the *kdpDE* genes were constructed (pYUB412_promoter insertion G - *whiB6* WT, pYUB412_promoter WT - *whiB6* WT, pYUB412_promoter WT - *whiB6* T51P, pYUB412_*kdpDE* WT, and pYUB412_*kdpDE* ΔCA). To obtain the *whiB6* and the *kdpDE* plasmids except for pYUB412_*kdpDE* ΔCA, the *whiB6* and the *kdpDE* genes and their natural promoter region were amplified by PCR using modified primers (Table S4), with additional EcoRV and AseI (or XbaI only for *kdpDE*) restriction sequences in the amplified fragment. The resulting PCR product was digested and ligated into the EcoRV-AseI-digested pYUB412 (or XbaI-digested pYUB412 for *kdpDE*). A mutagenesis of the pYUB412_*kdpDE* WT plasmid was performed to obtain the pYUB412_*kdpDE* ΔCA plasmid using the QuickChange II XL Site-Directed Mutagenesis Kit (Agilent Technologies). The pYUB412-based complementation plasmids were used to transform the *M. tuberculosis* H37RvΔ*whiB6* and H37RvΔ*kdpDE* mutant strains. Transformed clones were selected on solid medium for resistance to zeomycin and hygromycin.

### Strain collection

The study comprises *M. tuberculosis* strains received at the French National Reference Centre of Mycobacteria from French clinical laboratories most of the time for suspicion of resistant TB. Ethical review and approval were not required for this study in accordance with the local legislation and institutional requirements. Overall, 489 *M. tuberculosis* isolates were analyzed, including 32 isolates of which 3 non MDR with MIRU-VNTR genotype MLVA Mtbc 15-9 “100-32”, which has been linked with strains previously designated as “the successful Russian clone” (12). Isolates were sampled between 2017 and 2021. All MDR 100-32 strains were resistant to isoniazid due to the well-known drug resistance mutation S315T in KatG. Rates of resistance were high for second-line drugs, *i.e.*, fluoroquinolones (41%) and aminosides (55%).

### DNA isolation, library preparation and sequencing

*M. tuberculosis* genomic DNA was extracted using the GeneLEAD VIII system (Diagenode). Libraries were prepared with the Nextera XT DNA Library Preparation kit (Illumina) and sequenced on the Illumina NextSeq 500 at the Mutualized Platform for Microbiology (P2M, Institut Pasteur). Raw fastq data were uploaded into BioNumerics software vx (Applied Maths), which mapped the reads against the reference sequence (*M. tuberculosis* H37Rv, GenBank accession number NC_000962.3) and detected the single-nucleotide polymorphisms (SNPs). For greater accuracy, strict SNP filtering that removed positions in PE-PPE-PGRS genes was applied. The retained SNP positions had a minimum of 5x coverage and the minimum distance between SNPs was at least 12 base pairs (bp). All SNPs were manually checked by visualizing the corresponding read alignments. The fastq files were also uploaded into Phyresse (42). Only SNPs detected by both tools were retained. Functional annotation and categorization of affected genes was retrieved from the MycoBrowser (43).

### *In vitro* growth curves

All strains were diluted to an initial OD_600_ of 0.05 in Middlebrook 7H9 medium, and the OD_600_ was recorded at different time points over a period of 9 days. Data from three independent experiments were used for growth representation.

### RNA isolation, library preparation and sequencing

Mycobacterial strains were pelleted and bead beated in 1 mL of TRIzol (Life Technologies) with 0.1 mm silica beads (MP Biomedicals). After centrifugation, supernatants were extracted with chloroform, and RNA was precipitated with isopropanol and glycogen. RNA pellets were washed with 75% ethanol and dissolved in RNase-free water. Contaminant DNA was removed by incubation with DNase (TURBO DNA-*free*^TM^ kit, Life Technologies). RNA cleanup was performed with the RNeasy® Mini Kit (Qiagen). Three independent cultures for each strain were used for this experiment. Total RNA concentration was measured using the Qubit RNA HS assay kit. The quality of all samples was assessed with an Agilent Bioanalyzer device (Agilent Technologies) to verify RNA integrity. RNA-seq libraries were prepared using the Stranded Total RNA Prep and Ligation with Ribo-Zero Plus kit (Illumina) and sequenced using a NextSeq 2000 device (Illumina). Generated strand-specific 50-bp single-end reads were mapped against the reference genome of *M. tuberculosis* H37Rv (GenBank accession number AL123456.3) (44) using BWA-MEM v0.7.17-r1188 (45) (parameters: -M; -h 1000). Uniquely-mapped reads were extracted from the alignment maps according to the XA tag using the Python wrapper pysam v0.20.0 (https://github.com/pysam-developers/pysam) of SAMtools (46). Reads mapped on gene features were counted using featureCounts v2.0.4 (47) (parameters: -s 2; --primary). Counts associated with the genes *rrs*, *rrl* and *rrf*, encoding ribosomal RNAs, were excluded to prevent differences related to variable ribodepletion efficiencies during the library preparation of the samples. Gene counts were normalized and transformed by regularized logarithm using DESeq2 v1.38.3 (48). Exploration of unwanted variation within RNA-seq data using the R package sva v3.46.0 (49) revealed the presence of a batch effect between the first set of replicates and both second and third sets of replicates. Differential expression analysis was then performed using DESeq2 v1.38.3 (48) with a false-discovery rate (FDR, alpha) of 0.05 and the design formula ∼Condition + Batch, where Condition distinguished the different genotypes and growth conditions, and Batch allowed to take into consideration the batch effect detected with the surrogate variable analysis. Genes with an adjusted p-value (padj) lower than 0.05 were considered as differentially expressed (DE). Intersections of DE genes were computed using the R package UpSetR v1.4.0 (50). Functional annotation and categorization of DE genes was retrieved from the MycoBrowser (43). For visualization of gene expression levels, normalized and transformed gene counts were first corrected using the removeBatchEffect function from the R package limma v3.54.2 (51) and then ploted as heat maps using the R package pheatmap v1.0.12 (https://cran.r-project.org/package=pheatmap). Hierarchical clustering of genes was performed using the complete-linkage method on Pearson correlation distances.

### Secretion analysis and immunoblotting

Secretion analysis of *M. tuberculosis* was performed as described before (52). Briefly, bacteria were cultured until mid-logarithmic phase. Culture supernatants were recovered and proteins were precipitated with 10% (w/v) TCA. To obtain total lysates, mycobacterial pellets were washed twice and resuspended in PBS. Bacterial cells were broken by shaking with 0.1 mm silica beads (MP Biomedicals) for 8 min in a Tissue Lyser apparatus (Qiagen). Suspensions were centrifuged at 1,000 g for 3 min and the supernatant fraction obtained represented the total-cell lysate. Immunoblot analyses were performed using anti-EsxA antibodies (Hyb76-8, Antibodyshop, Statens Serum Institut) (53), anti-EsxB antibodies (a kind gift from I. Rosenkrands, Statens Serum Institut, Copenhagen, Denmark), and anti-SigA antibodies (a kind gift from I. Rosenkrands, Statens Serum Institut, Copenhagen, Denmark).

### *In vivo* virulence assessment of *M. tuberculosis* strains in a BALB/c mouse model of pulmonary tuberculosis

The experimental project was evaluated by the ethics committee n°005 Charles Darwin localized at the Pitié-Salpêtrière Hospital and approved by the French Ministry of Higher Education and Research under the number APAFIS#20300-2019041811145911 v3. Our animal facility received the authorization to carry out animal experiments (license number D75-13-08). The persons who carried out the animal experiments had followed a specific training recognized by the French Ministry of Higher Education and Research and follow the European and the French recommendations on the continuous training.

Mice infection experiment was performed as previously described (54). Briefly, six-week-old female BALB/cJRj mice (Janvier Labs, Le Genest Saint Isle, France) were intravenously infected in the tail with 0.5 mL of a bacterial suspension of one of the 8 following *M. tuberculosis* strains, H37Rv (1.5×10^7^ CFU), the Δ*whiB6* mutant strain (1.12×10^7^ CFU), the promoter G insertion - *whiB6* WT (2.25×10^7^ CFU), the promoter WT - *whiB6* WT (10^7^ CFU), and the promoter WT - *whiB6* T51P complemented strains (5×10^7^ CFU), the Δ*kdpDE* mutant strain (2×10^7^ CFU), the *kdpDE* WT (4.9×10^7^ CFU) and ΔCA (1.9×10^7^ CFU) complemented strains. Each group contained 20 mice which were randomly allocated in 3 groups of observation: 5 mice were euthanasied one day after infection, 5 after one month, and 10 after two months. The severity of infection was assessed by lung CFU counts. Lungs were aseptically removed and then homogenized with a GentleMacs Octo Dissociator (Miltenyi) in a volume of 2 mL of sterile distilled water. The number of CFU was then determined by plating homogenized lung suspensions in triplicate on Middlebrook 7H11 medium supplemented with 10% OADC and possibly with antibiotic. The CFU count was assessed after 6 weeks of incubation at 37°C.

### Statistical analyses

Potential statistical differences in bacterial loads were evaluated by Kruskal-Wallis test using GraphPad Prism 5.

## Acknowledgements

Support by grants from the Agence National de Recherche (ANR-10-LABX-62-IBEID and ANR-20-CE15-0013) are gratefully acknowledged. The Biomics Platform is supported by France Génomique (ANR-10-INBS-09) and IBISA. The French National Reference Centre of Mycobacteria is supported by an annual grant from Santé Publique France.

We thank Alexandre Pawlik for advice in transcriptome preparation, and Azimdine Habib, Laure Lemée and Marc Monot (Biomics Platform, C2RT, Institut Pasteur, Paris, France) for processing the RNA sequencing.

## Data availability statement

Whole-genome sequencing data have been deposited in the Sequence Read Archive (SRA) under the BioProject accession number PRJNA1091467. RNA-seq data have been deposited in the Gene Expression Omnibus (GEO) database under the SuperSeries accession number GSE269919.

## Supporting information

**Table S1.** Transcriptional changes associated with the *whiB6* genotypes relative to H37Rv.

**Table S2.** Transcriptional changes associated with the *kdpDE* genotypes relative to H37Rv in the presence of potassium.

**Table S3.** Transcriptional changes associated with the *kdpDE* genotypes relative to H37Rv in the absence of potassium.

**Table S4.** Primers sequences.

**Fig. S1.** Functional annotation and categorization of genes detected as differentially expressed in the *whiB6* genotypes relative to H37Rv.

**Fig. S2.** Functional annotation and categorization of genes detected as differentially expressed in the *kdpDE* genotypes relative to H37Rv in the presence of potassium.

**Fig. S3.** Functional annotation and categorization of genes detected as differentially expressed in the *kdpDE* genotypes relative to H37Rv in the absence of potassium.

**Fig. S4.** Virulence evaluation of *whiB6* T51P and *kdpDE* ΔCA mutations in BALB/c mice.

