## Supplementary Figures for "Evaluation of the role of *whiB6* and *kdpDE* in the dominant multidrug resistant clone *Mycobacterium tuberculosis* B0/W148"

**Fig. S1. Functional annotation and categorization of genes detected as differentially expressed in the *whiB6* genotypes relative to H37Rv.** Gene overlap is depicted as a proportion of differentially expressed genes belonging to each functional category relative to the total number of genes detected as differentially expressed in each *whiB6* genotype as compared to H37Rv (x-axis, n). Total number of genes annotated within each functional category is indicated (y-axis, N).

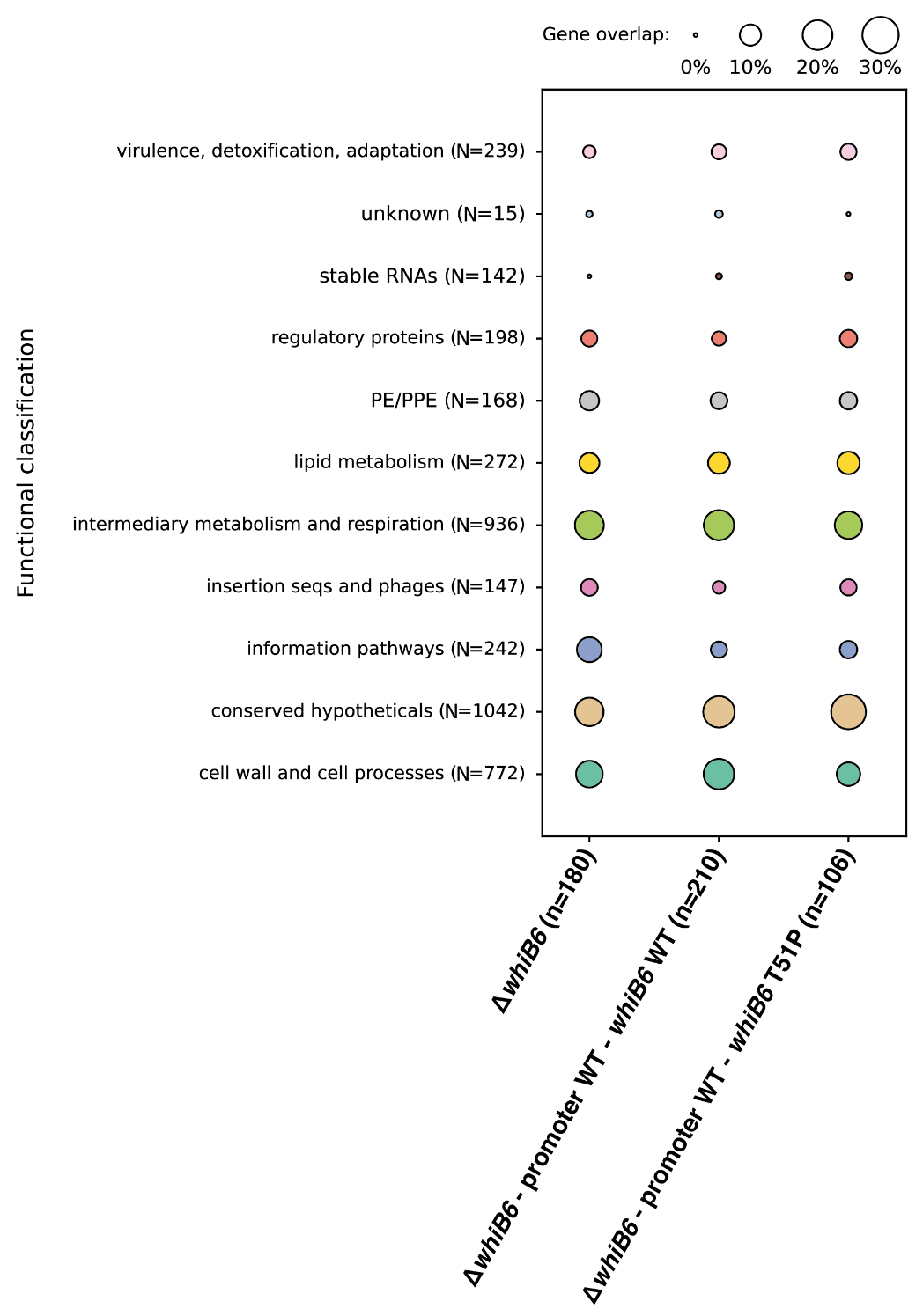

**Fig. S2. Functional annotation and categorization of genes detected as differentially expressed in the *kdpDE* genotypes relative to H37Rv in the presence of potassium.** Gene overlap is depicted as a proportion of differentially expressed genes belonging to each functional category relative to the total number of genes detected as differentially expressed in each *kdpDE* genotype as compared to H37Rv during normal growth condition (x-axis, n). Total number of genes annotated within each functional category is indicated (y-axis, N).

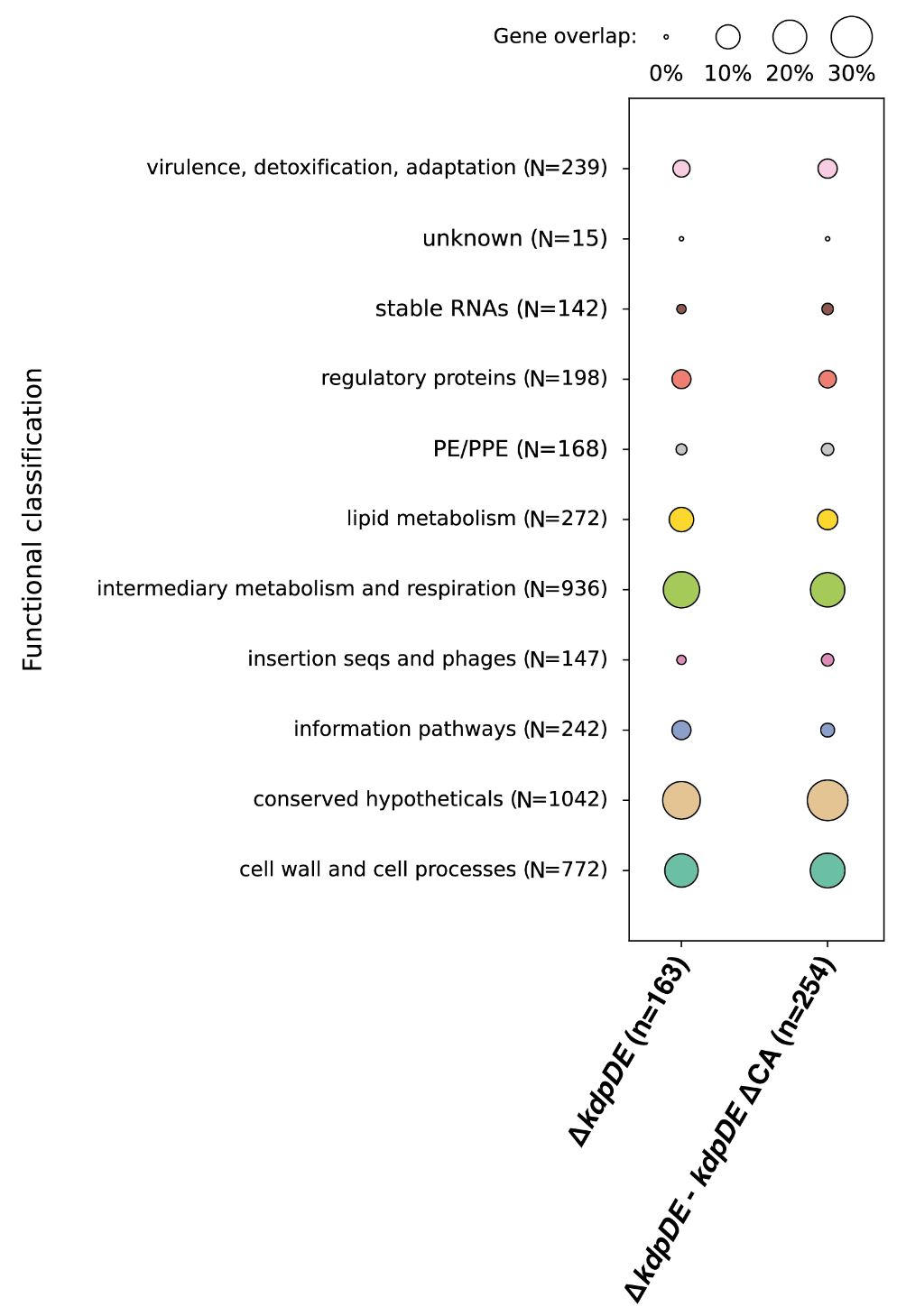

**Fig. S3. Functional annotation and categorization of genes detected as differentially expressed in the *kdpDE* genotypes relative to H37Rv in the presence of potassium.** Gene overlap is depicted as a proportion of differentially expressed genes belonging to each functional category relative to the total number of genes detected as differentially expressed in each *kdpDE* genotype as compared to H37Rv upon potassium depletion (x-axis, n). Total number of genes annotated within each functional category is indicated (y-axis, N).

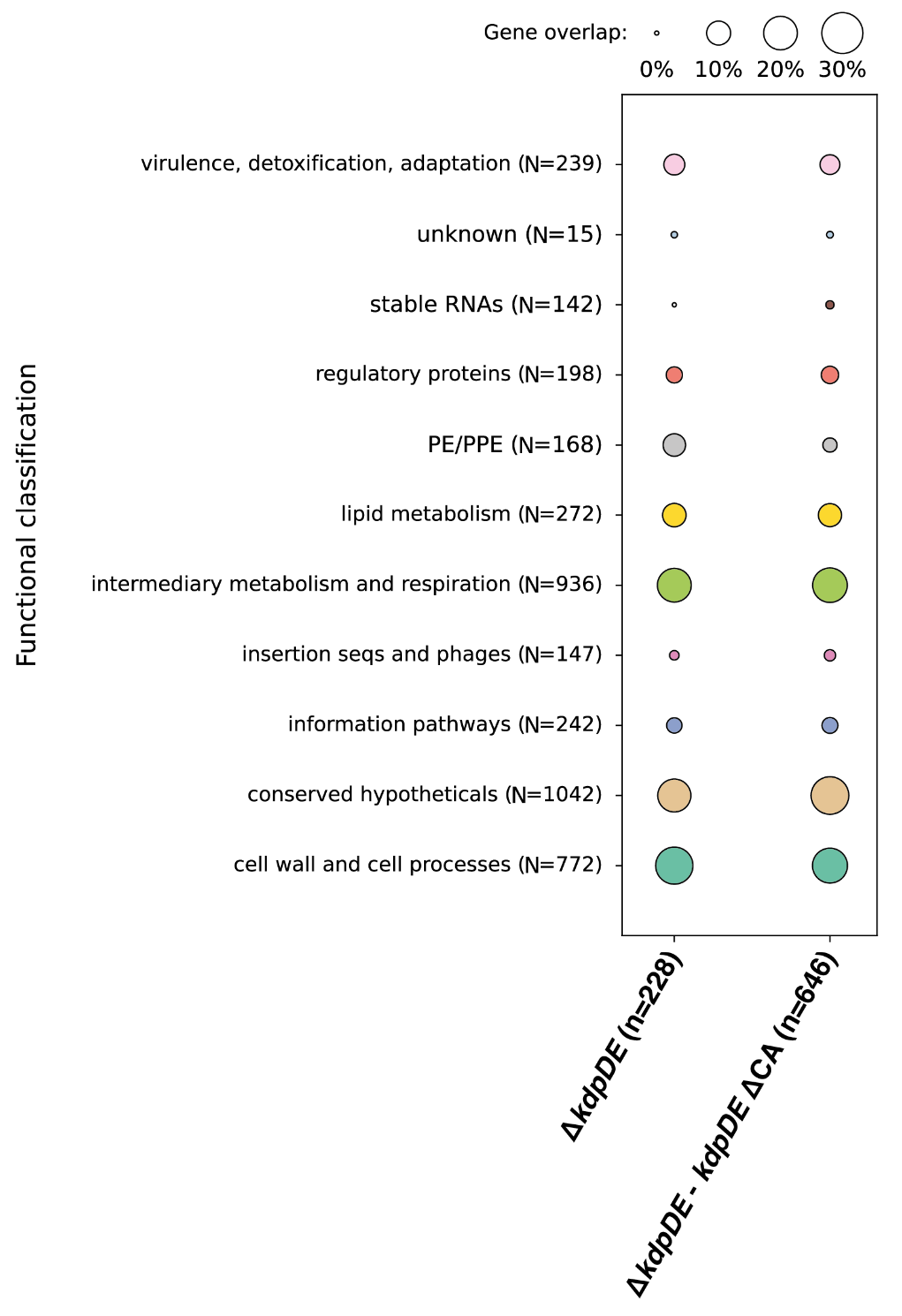

**Fig. S4. Virulence evaluation of *whiB6* T51P and *kdpDE* ΔCA mutations in BALB/c mice.** (A) Time course after mouse infection with H37Rv, the ∆*whiB6* mutant strain, and the promoter G insertion - *whiB6* WT, the promoter WT - *whiB6* WT, and the promoter WT - *whiB6* T51P complemented strains. (B) Time course after mouse infection with H37Rv, the ∆*kdpDE* mutant strain, and both *kdpDE* WT and ∆CA complemented strains. Recovery of the bacteria is enumerated by CFU per lung at 1, 30, 60 days after injection. Means for 5 mice at day 1 and 30 or for 10 mice at day 60 are shown with one point representing a mouse. Extra points correspond to mice killed between day 30 and day 60 due to probable otitis; they are attributed to day 30 or day 60 depending on the closest date.

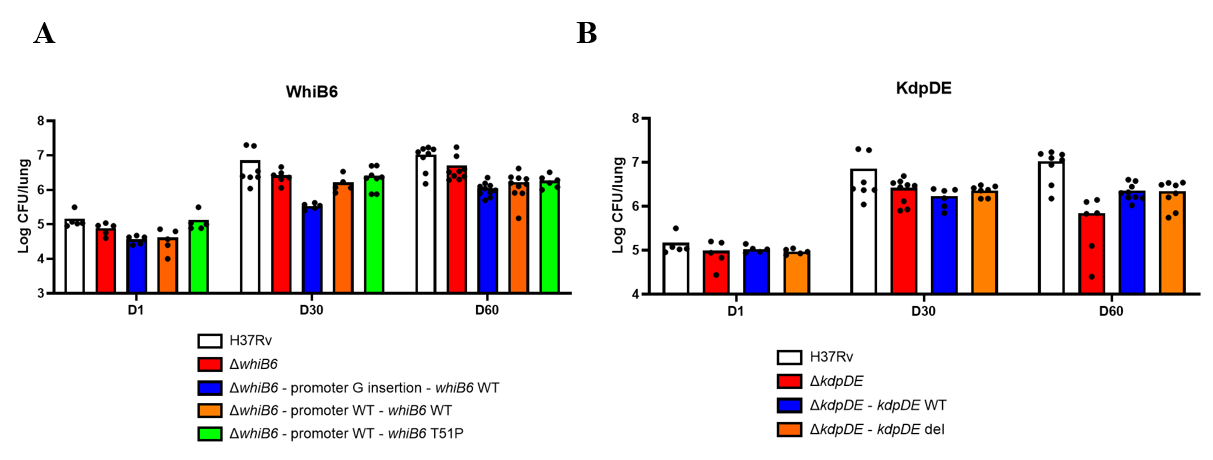

**Table S4. Primers sequences.**

|  | **Name** | **Sequence** |
| --- | --- | --- |
|  | Construction of Mtb H37Rv∆*whiB6* | |
| **1** | Up-region_whiB6_Forward | 5’-GATTCGGACGGACACGCCGA-3’ |
| **2** | Up-region_whiB6_Reverse | 5’-ttcgttttatttgatgcctgTCAGGCCGGGCGCGGGCATT-3’ |
| **3** | Down-region_whiB6_Forward | 5’-agttttcgttccactgagcgTAACCATAGCGATGCAACAG-3’ |
| **4** | Down-region_whiB6_Reverse | 5’-CCAATGATCGCGACACCGCG-3’ |
|  | Construction of pYUB412_promoter insertion G - *whiB6* WT, pYUB412_promoter WT - *whiB6* WT, pYUB412_promoter WT - *whiB6* T51P | |
| **5** | WhiB6_EcoRV_Forward | 5’-GTAGGATATCGACGCCTAACCGTTGCACCCTTCT-3’ |
| **6** | WhiB6_AseI_Reverse | 5’-CGGCATTAATCTTCTCATGCCGATTGGGCAGACA-3’ |
|  | Construction of Mtb H37Rv∆*kdpDE* | |
| **7** | Up-region_kdpDE_del_Forward | 5’-cgtctaagaaaccTGAAGGCGACATGGTCGG-3’ |
| **8** | Up-region_kdpDE_del_Reverse | 5’-tcgactgagccttGTGCACAAGAATCGAGAGG-3’ |
| **9** | Down-region_kdpDE_del_Forward | 5’-cctgcatgaccaaTTTCCGCTGGGAGCGGATG-3’ |
| **10** | Down-region_kdpDE_del_Reverse | 5’-gacttcagagcttTCGTGGTCGAAGAGTTGGC-3’ |
|  | Construction of pYUB412_*kdpDE* WT plasmid | |
| **11** | KdpDE_XbaI_Forward | 5’-ccctctagaGTTGTCGACCGTAGTCAT-3’ |
| **12** | KdpDE_XbaI_Reverse | 5’-ccctctagaGTCACATCTGCCACACG-3’ |
|  | Construction of pYUB412_*kdpDE* del plasmid | |
| **13** | KdpDE_pYUB412_Forward | 5’-GGGCGGCGGGCTCAGTGGTGATCG-3’ |
| **14** | KdpDE_pYUB412_Reverse | 5’-CGATCACCACTGAGCCCGCCGCCC-3’ |
